# Molecular and functional properties of human *Plasmodium falciparum* CSP C-terminus antibodies

**DOI:** 10.1101/2023.01.19.524692

**Authors:** Opeyemi Ernest Oludada, Giulia Costa, Clare Burn Aschner, Anna S. Obraztsova, Katherine Prieto, Caterina Canetta, Stephen L. Hoffman, Peter G. Kremsner, Benjamin Mordmüller, Rajagopal Murugan, Jean-Philippe Julien, Elena A. Levashina, Hedda Wardemann

**Author notes:** equal contribution (Rajagopal Murugan, Jean-Philippe Julien, Elena A. Levashina, Hedda Wardemann). Correspondence: Hedda Wardemann, Elena A. Levashina, Jean-Philippe Julien.

## Abstract

Human monoclonal antibodies (mAbs) against the central repeat and junction domain of *Plasmodium falciparum* circumsporozoite protein (PfCSP) have been studied extensively to guide malaria vaccine design compared to antibodies against the PfCSP C terminus. Here, we describe the molecular characteristics and protective potential of a panel of 73 germline and mutated human mAbs against the highly immunogenic PfCSP C-terminal domain. Two mAbs recognized linear epitopes in the C-terminal linker with sequence similarity to repeat and junction motifs, whereas all others targeted conformational epitopes in the α-thrombospondin repeat (α-TSR) domain. Specificity for the polymorphic Th2R/Th3R but not the conserved RII+ region in the α-TSR was associated with *IGHV3-21*/*IGVL3-21* or *IGLV3-1* gene usage. Although the C terminus specific mAbs showed signs of more efficient affinity maturation and class-switching compared to anti-repeat mAbs, parasite inhibitory activity was limited to a single C-linker reactive mAb with cross-reactivity to the central repeat and junction. The data provide novel insights in the human anti-C-linker and anti-α-TSR antibody response that support exclusion of the PfCSP C terminus from malaria vaccine designs.

## INTRODUCTION

*Plasmodium falciparum* (Pf) malaria is a mosquito-borne parasitic disease that is highly endemic in sub-Saharan Africa, where it remains the primary cause of childhood illness and death (WHO, 2021). RTS,S/AS01 (Mosquirix™), the only marketed malaria vaccine, has been recommended for wide-spread use among children in areas with high to moderate Pf transmission.

RTS,S/AS01 is a subunit vaccine that targets Pf circumsporozoite protein (PfCSP), the major protein on the surface of sporozoites that are transmitted to humans during the blood meal of infected *Anopheles* mosquitoes (Swearingen *et al*, 2016). PfCSP consists of three domains, a poorly characterized N-terminal domain (N-CSP), an unstructured central repeat region with large numbers of repeating four amino-acid motifs, and a C-terminal domain (C-CSP) with a linker (C-linker) and α-thrombospondin type-1 repeat (α-TSR) subdomain that is attached to the cell membrane by a GPI anchor (Dame *et al*, 1984; Doud *et al*, 2012; McCutchan *et al*, 1996; Wang *et al*, 2005). RTS,S/AS01 contains a genetic fusion protein of an N-terminally truncated form of PfCSP from the Pf clone 3D7 derived from the laboratory strain NF54 and hepatitis B surface antigen, which is complexed with free HbsAg to form virus-like particles and boost immunogenicity (Bojang *et al*, 2001; Casares *et al*, 2010; Gordon *et al*, 1995; RTS *et al*, 2011; Stoute *et al*, 1997). The PfCSP part of RTS,S comprises 18.5 of the 38 repeating asparagine (N) – alanine (A) - asparagine (N) – proline (P) motifs in the central repeat domain (Stoute *et al*., 1997; Zavala *et al*, 1985) and the complete C-CSP. The characteristic NANP motifs are fully conserved in Pf parasites (with only slight variations in the number of repeating units) independent of their geographic origin and are long known to be the target epitopes of protective antibodies (Nardin *et al*, 1982; Zavala *et al*, 1983).

The successful clinical development of RTS,S/AS01 in contrast to the other numerous malaria vaccine candidates confirms the potency of PfCSP as a promising vaccine target. However, the overall efficacy is limited and protection is short-lived (Agnandji *et al*, 2011; Asante *et al*, 2011; Olotu *et al*, 2016; Olotu *et al*, 2011). To guide the design of a more efficacious PfCSP-based malaria vaccine, recent studies have characterized large numbers of human monoclonal antibodies (mAbs) against PfCSP (Kisalu *et al*, 2018; Murugan *et al*, 2018; Oyen *et al*, 2017; Tan *et al*, 2018; Triller *et al*, 2017; Wang *et al*, 2020). Their molecular and functional characterization provided deep insights in the development and potency of anti-repeat antibodies and identified new epitopes of protective mAbs in the N-terminal junction with alternating NANP and NANP-like (NPDP, NVDP) motifs that is not contained in RTS,S.

In contrast to the large numbers of well-characterized antibodies against the central repeat domain and N-terminal junction, the role of antibodies against the N and C terminus remains controversial. The PfCSP N terminus is overall poorly immunogenic and undergoes proteolytic cleavage at region I (RI), a stretch of six amino acids that is highly conserved in all *Plasmodium* species (Coppi *et al*, 2005; Espinosa *et al*, 2015). Human monoclonal antibodies against the N terminus distal to RI have not been described, and the few known mouse monoclonal antibodies show little sporozoite reactivity and parasite inhibitory activity (Herrera *et al*, 2015; Thai *et al*, 2020).

Although immunization with recombinant PfCSP induces dominant humoral responses against the C-CSP in animal models and humans (Cawlfield *et al*, 2019; Genito *et al*, 2017; Hutter *et al*, 2022), the limited number of human C-CSP mAbs that have been described so far did not allow deep analyses of their gene features, epitope breadth or preferences and the parasite-inhibitory activity. Two target sites of human monoclonal antibodies have been identified in the highly polymorphic α-TSR domain and the conserved RII+ region (Beutler *et al*, 2022; Scally *et al*, 2018). However, whether binding to these epitopes is associated with specific Ig genes, whether other epitopes are targeted by anti-C-CSP antibodies and whether epitope specificity is linked to differences in parasite inhibition remains unclear. To address these questions, we characterized the molecular, functional properties, and binding-specificity of a large panel of recombinant human monoclonal antibodies cloned from memory B cells of malaria-naïve individuals immunized with aseptic, cryopreserved, radiation-attenuated Pf sporozoites (PfSPZ Vaccine; (Mordmuller *et al*, 2022)).

## RESULTS

### Immunization with radiation-attenuated *Plasmodium falciparum* sporozoites (PfSPZ) induces antibody responses against C-CSP

To identify individuals with antibody responses against C-CSP, we measured the humoral response in sera from 12 malaria-naive European volunteers 28 days after three direct venous inoculation (DVI) immunizations with 900,000 aseptic, purified, radiation-attenuated Pf NF54 sporozoites (Sanaria® PfSPZ Vaccine; (Mordmuller *et al*., 2022)). Comparison of the IgG response against a peptide representing the central NANP repeat (NANP_5_) and against C-CSP showed responses to both domains (Fig. 1A; Fig. S1A; Table S1).

**Figure 1:**
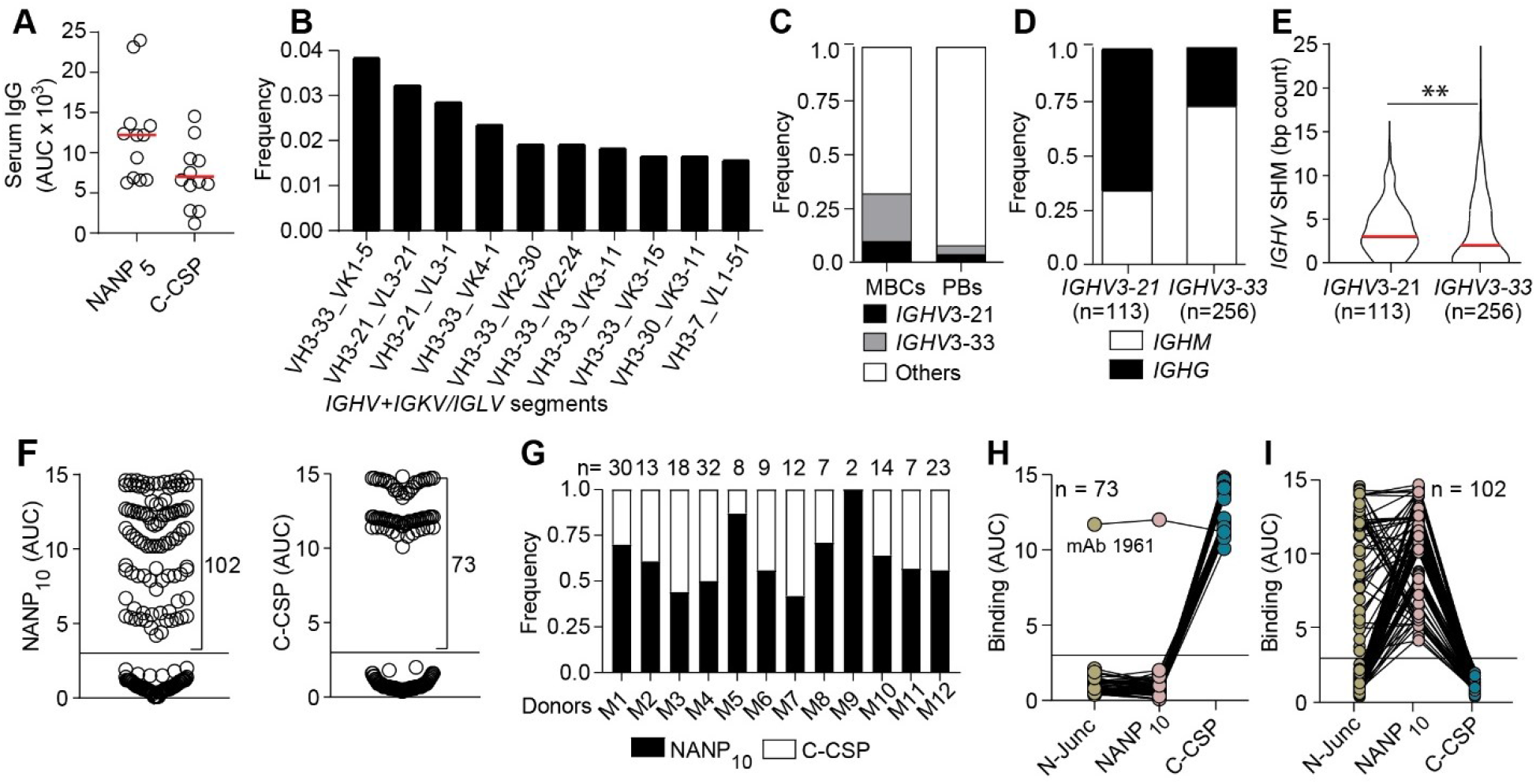
B cell response against the PfCSP C terminus. **(A)** Anti-NANP_5_ and anti-C-CSP serum IgG. Circles represent ELISA area under the curve (AUC) values for individual donors (n=12). **(B)** Ten most frequent Ig heavy and light chain V gene pairs in PfCSP-reactive memory B cells (MBCs; n=1172). **(C)** Usage frequency of *IGHV3-21, IGHV3-33* and other *IGHV* genes in PfCSP-reactive MBCs (n=1172) and plasmablasts (PBs; n=2380). **(D and E)** Isotype distribution and somatic hypermutation (SHM) count **(E)** in *IGHV3-21* and *IGHV3-33* genes from MBCs in (B, C). **(F)** NANP_10_ (left) and C-CSP (right) binding of FL-CSP-reactive mAbs (n=177). Circles represent ELISA AUC values for each mAb. **(G)** Frequency of NANP_10_ and C-CSP reactive mAbs per donor. The number of tested mAbs is indicated. **(H and I)** Cross-reactivity of C-CSP **(H)** and NANP_10_ **(I)** reactive mAbs with the PfCSP N-junc, NANP_10_ and C-CSP. Red lines in **(E)** indicate mean values. Black horizontal lines in **(F), (H)** and **(I)** indicate the threshold for binding. \*\**P* < 0.01, two-tailed Mann-Whitney test **(E)**. Data in **(A)** are representative of two independent experiments. Data in **F, H** and **I** data show means from three independent experiments.

To directly compare the anti-C-CSP and anti-repeat antibody response at monoclonal level, we isolated single circulating PfCSP-reactive memory B cells from all donors 14 and 35 days after the third immunization (III+14, III+35; Fig. S1B and S1C) and amplified the paired immunoglobulin (Ig) heavy and light chain genes. Sequencing of the RT-PCR Ig gene products showed that in all donors at both time points, the anti-PfCSP response was dominated by memory B cells expressing VH3-21 and VH3-33 antibodies, frequently in association with Vλ3-21 or Vλ3-1 and Vκ1-5 light chains, respectively (Fig. 1B; Fig. S1D). Both *IGHV* genes were enriched in the PfCSP-reactive memory B cell pool compared to circulating plasmablasts from the same individuals and time points that represent the B cell response to all sporozoite antigens (Fig. 1C). The high abundance of VH3-33 expressing PfCSP-reactive memory B cells and frequent association with Vκ1-5 suggested that these cells recognized the central repeat, similar to VH3-33 antibodies induced by immunization with non-attenuated sporozoites (Murugan *et al*., 2018). Compared to VH3-33 antibodies, which were mostly IgM carrying few somatic hypermutations (SHM), the VH3-21 response was dominated by IgG cells with higher mean SHM counts (Fig. 1D and 1E), indicating their participation in germinal center (GC) responses.

To determine how VH3-21 usage was linked to PfCSP-reactivity, we cloned and expressed 267 monoclonal antibodies (mAbs) covering the overall repertoire diversity of PfCSP-reactive memory B cells, including 42 VH3-21 and 47 VH3-33 antibodies with similar isotype distribution (Fig. S1E; Table S2 and S3). From this panel, 177 (66%) recombinant mAbs showed ELISA reactivity with full-length PfCSP (FL-CSP) and lacked binding to non-related antigens, demonstrating their specificity for PfCSP (Fig. S1F and G). Of these FL-CSP specific mAbs, 102 showed reactivity to a peptide representing the central NANP repeat (NANP_10_), and 73 mAbs recognized the C-CSP domain (Fig. 1F). Two mAbs lacked binding to NANP_10_ or C-CSP, but bound a peptide covering the junction motifs (N-junc) (Fig. S1H). NANP-reactive mAbs were identified in all donors and C-CSP-reactive mAbs in all but one donor (Fig. 1G).

To determine whether any of the C-CSP-reactive antibodies showed cross-reactivity with the repeat domain including the N-terminal junction, as previously reported (Murugan *et al*, 2020), we measured binding of the mAbs to NANP_10_ and the N-junc peptide. With one exception (mAb 1961), all C-CSP-reactive mAbs were highly specific, whereas most of the NANP_10_-binders showed cross-reactivity to the junction as expected (Fig. 1H and1I). Thus, immunization with radiation-attenuated sporozoites elicited PfCSP C-CSP specific and repeat/junction cross-reactive humoral and memory B cell responses associated with *IGHV3-21* and *IGHV3-33* gene signatures, respectively.

### C-CSP specific mAbs are frequently encoded by *IGHV3-21*

To further characterize the panel of 73 C-CSP specific mAbs, we determined their binding strength by surface plasmon resonance (SPR; Fig. 2A). The mAbs recognized C-CSP with a wide range of affinities (10^−5^ to 10^−10^ M). Ig gene sequence analysis showed that half (37/73) of all C-CSP binders were encoded by *IGHV3-21* most of which paired with *IGLV3-21* or *IGLV1-3* genes, demonstrating the strong link between this gene combination and anti-C-CSP antibody reactivity (Fig. 2B; Fig. S2A). The other half of the anti-C-CSP antibodies used diverse gene combinations and showed no significant difference in affinity, isotype distribution or somatic mutation load compared to VH3-21 mAbs (Fig. 2B; Fig. S2B-2D). As expected, VH3-33 heavy chains paired with Vκ1-5 light chains were highly abundant among the NANP-reactive mAbs, similar to VH3-33/Vκ1-5 anti-repeat antibodies induced by immunization with non-irradiated sporozoites (Fig. S2A and S2E).

**Figure 2:**
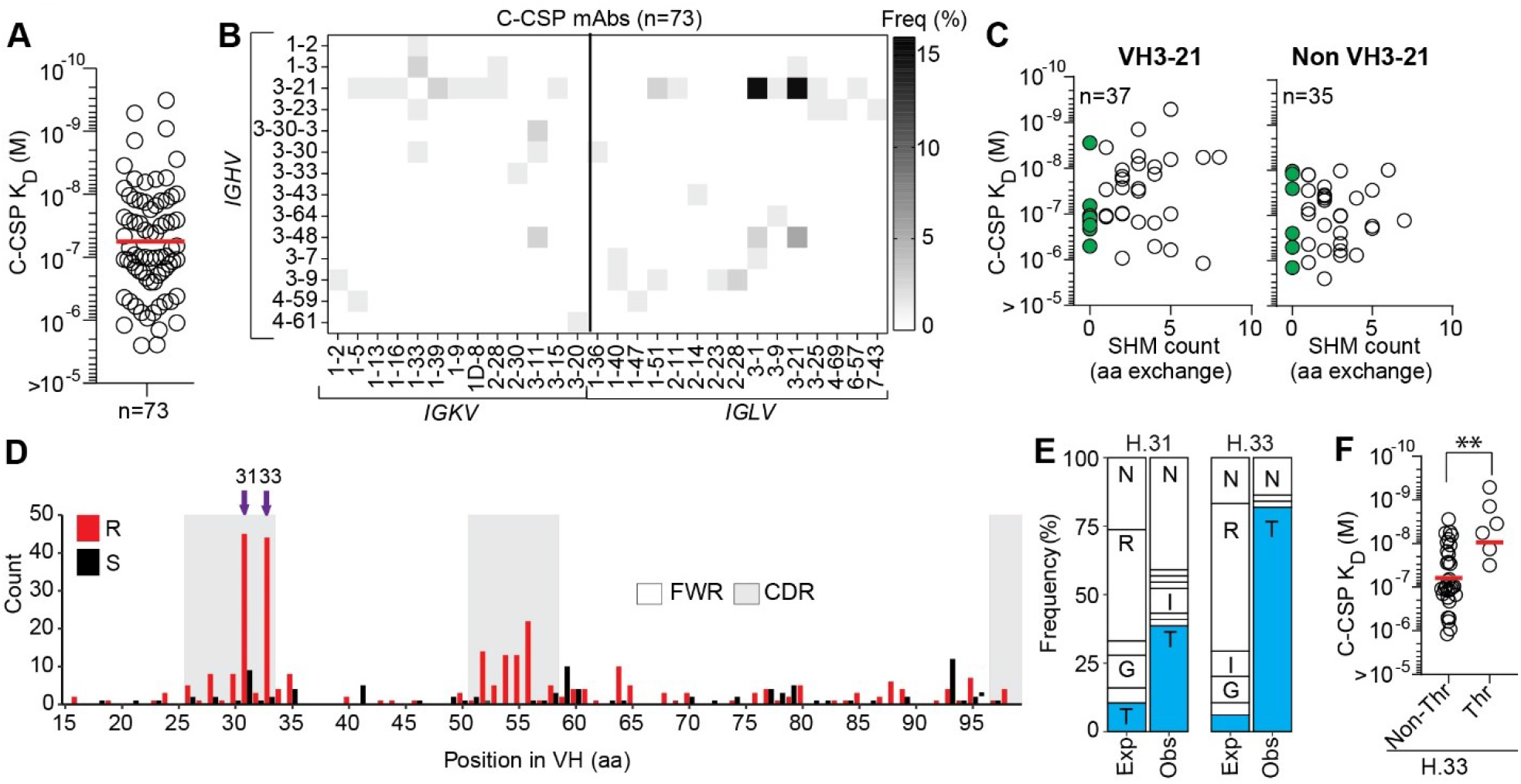
C-CSP-specific mAbs are frequently encoded by *IGHV3-21*. **(A)** SPR affinity. **(B)** Frequency of mAbs encoded by the indicated *IGHV* and *IGKV* or *IGLV* pairs. **(C)** VH SHM load of VH3-21 and non-VH3-21 mAbs. mAbs with unmutated VH are highlighted in green. **(D)** Amino acid (aa) VH replacement (red bars) and silent (black bars) SHM in VH3-21 mAbs (n=113). FWR, framework region; CDR, complementarity-determining region. **(E)** Observed (Obs) aa usage frequency at position H.31 and H.33 in VH3-21 mAbs carrying a replacement mutation at these positions compared to the expected (Exp) neutral mutation model (Gupta *et al*, 2015; Yaari *et al*, 2013). Single-letters indicate aa residues: G, Gly; I, Ile; N, Asn; R, Arg; T, Thr. **(F)** C-CSP affinity for selected VH3-21 mAbs with or without Thr mutation at position H.33. Red horizontal lines in **(A, F)** indicate mean values. \*\**P* < 0.01, two-tailed Mann-Whitney test **(F)**. Data in **A, C and F** are representative of two independent experiments.

Although most C-CSP-specific antibodies were class-switched and carried somatic mutations as signs of affinity maturation, we identified several VH3-21 and non-VH3-21 germline antibodies with high C-CSP affinity suggesting that the naïve human B cell repertoire contains numerous C-CSP-reactive precursor cells (Fig. 2C). Sequence alignments of all VH3-21 antibodies showed strong signs of selection indicated by enrichment of replacement mutations in CDRs compared to framework regions (FWR; Fig. 2D). Almost half of the VH3-21 antibodies carried replacement mutations at two positions in HCDR1 (H.31, H.33) and threonine was strongly enriched in these positions compared to the neutral mutation model (Fig. 2D and E; Fig S2F and G; (Gupta *et al*, 2015; Yaari *et al*, 2013)). VH3-21 mAbs carrying the selected H.S33T mutation showed on average higher C-CSP affinities than mAbs with the germline residue H.S33 or with non-selected mutations demonstrating the role of the H.S33T exchange in affinity maturation (Fig. 2F). The data shows a strong link of VH3-21 gene segment usage with C-CSP reactivity and provides direct evidence for effective affinity maturation of the anti-C-CSP response similar to the selection of H.S31N and H.V50I mutations in VH3-33 antibodies (Fig. S2H and 2I), which mediate high affinity to repeating NANP motifs (Imkeller *et al*, 2018).

### C-CSP specific antibodies preferentially target two distinct conformational epitopes in the α-TSR domain

To determine which of the two C-terminal subregions (Fig. 3A) the C-CSP-specific and C-CSP cross-reactive antibodies recognized, we measured binding to the C-linker (aa 273-310) and the α-TSR domain (aa 311-384) by ELISA (Fig. 3B; Fig. S3A). With two exceptions that recognized the C-linker region (mAb 3764, and mAb 1961 with NANP_10_ and N-junc cross-reactivity; Fig. 1H), all mAbs showed specificity for the α-TSR domain.

**Figure 3:**
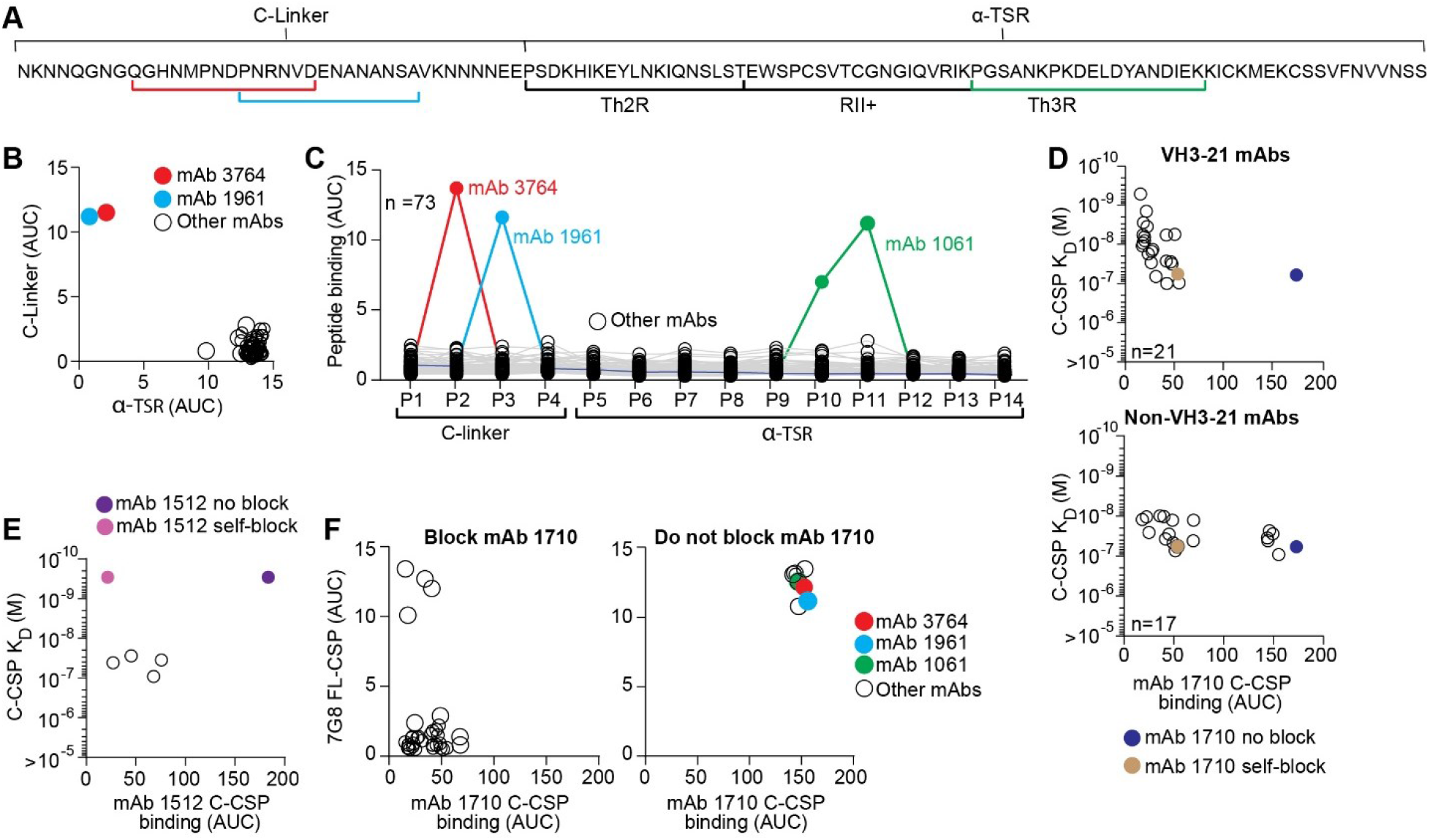
Epitope specificity of C-CSP-specific mAbs. **A**. NF54 PfCSP C-CSP aa sequence. The C-linker and α– TSR domain with Th2R, RII+ and Th3R are indicated. **(B and C)** ELISA reactivity of C-CSP-reactive mAbs (n=73) with the C-linker and α-TSR domain **(B)** and overlapping C-CSP peptides (P1-P14; Table S1 for sequences) **(C). (D)** C-CSP reactivity of the α-TSR-specific mAb 1710 (Scally *et al*, 2018) in a blocking ELISA with C-CSP-specific VH3-21 (n=21; upper panel) or non-VH3-21 (n=17; lower panel) mAbs with the indicated SPR affinities. mAb 1710 binding without blocking (blue) and after self-blocking (gold) is shown for comparison. C-CSP reactivity of mAb 1512 (Beutler *et al*, 2022) in a blocking ELISA with mAbs (with the indicated SPR affinities) that do not block C-CSP binding of mAb 1710 (open circles; D). mAb 1512 binding without blocking (purple) and after self-blocking (magenta) is shown for comparison. ELISA cross-reactivity with Pf-7G8 FL-CSP of CSP-specific mAbs that block (left) or do not block (right) mAb 1710 binding to NF54 C-CSP. Highlighted mAbs **(B, C** and **F)** with their binding sites (highlighted in **A)** are mAb 3764 (C-linker specific; red), mAb 1961 (C-linker, repeat and junction cross-reactive; light blue), mAb 1061 (α-TSR peptide 10 and peptide 11 specific; green).

To define whether any of the C-CSP-reactive antibodies recognized linear epitopes, we tested their binding to 14 overlapping peptides (P1-14) covering the complete NF54 C-CSP (Fig. 3C; Fig. S3B; Table S1). Only three mAbs showed reactivity in these assays: the C-linker specific mAb 3764 with specificity for P2 spanning aa 281-294 (QGHNMPNDPNRNVD) and the α-TSR-reactive mAb 1061, which bound P10 and P11 covering aa 345-366 in Th3R region. These two peptides overlap by six aa (KPKDEL), suggesting that the mAb 1061 target epitope covers aa KPKDEL. The C-CSP, junction and repeat cross-reactive mAb 1961 showed reactivity with P3 (PNRNVDENANANSA) suggesting that C-CSP binding of this mAb might be mediated via the NANA motif, similar to the previously reported C-CSP, junction and repeat cross-reactive mAb 3246 (Murugan *et al*., 2020). All other C-CSP specific mAbs (70/73) lacked peptide reactivity indicating that they recognized conformational epitopes.

Similar to the previously reported anti-α-TSR mAb 1710 (Scally *et al*., 2018), many of these antibodies were encoded by *IGHV3-21*, suggesting that these antibodies and mAb 1710 might recognize the same or overlapping epitopes in Th2R and Th3R. To address this question, we selected 21 *IGHV3-21*-encoded and 17 non-*IGHV3-21* encoded C-CSP specific mAbs with similar or higher C-CSP affinity compared to mAb 1710 (≤ 10^−7^ M) and tested their ability to block mAb 1710 binding to C-CSP by ELISA (Fig. 3D, Fig. S3C, Table S4). All 21 VH3-21 and most of the non-VH3-21 (12/17) mAbs blocked binding of mAb 1710 demonstrating that the vast majority of anti-C-CSP antibodies shared specificity for the same or overlapping epitopes in the Th2R/Th3R. Only five of the 17 non-VH3-21 mAbs with diverse Ig genes did not interfere with mAb 1710 binding (Fig. 3D). To determine their epitope specificity, we tested their ability to block mAb 1512, an RTS,S/AS01-induced C-CSP mAb with specificity for a conformational epitope in the conserved RII+ of the α-TSR subdomain (Beutler *et al*., 2022). All five antibodies blocked mAb 1512 binding, demonstrating their specificity for the conserved RII+ epitope independent of their Ig gene usage (Fig. 3E; Fig. S3D).

The Th2R and to lesser extent the C-linker and Th3R are highly polymorphic (Aragam *et al*, 2013; Gandhi *et al*, 2012). To evaluate whether the C-CSP-specific antibodies were able to accommodate these polymorphisms, we tested their ability to bind to PfCSP from 7G8, which differs from NF54 C-CSP by seven aa: one aa in the C-linker and six aa in the α-TSR region (five in Th2R and one in Th3R; Fig. S3E and F). With four exceptions, none of the mAb 1710-blocking mAbs recognized 7G8 PfCSP (Fig. 3E). In contrast, mAbs 3764, 1961 and 1061 with linear peptide epitopes in the C-linker and Th3R, respectively, and the five mAbs with reactivity to the conformational mAb 1512 epitope in RII+ showed cross-reactivity with 7G8 (Fig. 3F). Thus, only antibodies against the conserved linear or conformational C-CSP epitopes not overlapping with the polymorphic mAb 1710 binding site showed 7G8 cross-reactivity.

In summary, the vast majority of anti-C-CSP antibodies showed the same specificity as mAb 1710, demonstrating the strong immunogenicity of their polymorphic target epitopes in the α-TSR domain. Binding to this site was strongly linked with *IGHV3-21* gene usage since all VH3-21 mAbs bound this epitope. Antibodies against conserved epitopes in the α-TSR domain or the C-linker (conformational or linear) used diverse IgH and IgL combinations and showed cross-reactivity to PfCSP from 7G8, but were overall rare.

### C-CSP-specific antibodies fail to bind live sporozoites and lack parasite inhibitory activity

Differences in epitope specificity and Ig gene usage of antibodies might be linked to their parasite binding and inhibitory capacities. We selected 15 representative antibodies with high C-CSP affinity against the linear and conformational epitopes in the C-linker and α-TSR domain and measured their ability to bind live fluorescent *P. berghei* PfCSP(mCherry) sporozoites expressing PfCSP (Fig. S4A; Table S5). Compared to the anti-repeat mAb 2A10 (Triller *et al*., 2017), which showed strong binding at low concentration (1 µg/ml) in flow cytometry, none of the mAb 1710- or mAb 1512-like anti-α-TSR mAbs recognized live sporozoites even at 100 μg/ml regardless of their epitope specificity and affinity (Fig. 4A), supporting the earlier observations that the α-TSR domain on the surface of live sporozoites was not accessible for antibody binding (Scally *et al*., 2018; Wang *et al*, 2021). Weak binding was only observed for the C-linker specific mAb 3764 at high (100 µg/ml) but not at low (1 µg/ml) concentration. However, the cross-reactive mAb 1961 strongly bound to the sporozoites similar to the positive control anti-repeat mAb 2A10 (Fig. 4B-4D). In agreement with their inability to bind live sporozoites, none of the anti-C-CSP specific antibodies showed Pf sporozoite traversal-inhibitory capacity even at a high concentration (100 µg/ml), including the high-affinity C-linker specific mAb 3764, which weakly bound sporozoites (Fig. 4E). Only the repeat and junction cross-reactive mAb 1961 showed *in vitro* sporozoite-inhibitory activity in the range of mAbs 2A10 and 317, a high affinity human anti-NANP mAb (Oyen *et al*., 2017; Triller *et al*., 2017). Nevertheless, its capacity to protect from the development of parasitemia *in vivo* after passive transfer in mice was significantly lower than that of mAb 317 and similar to mAb 1710 (Fig. 4F; Fig. S4B; Table S6).

**Figure 4:**
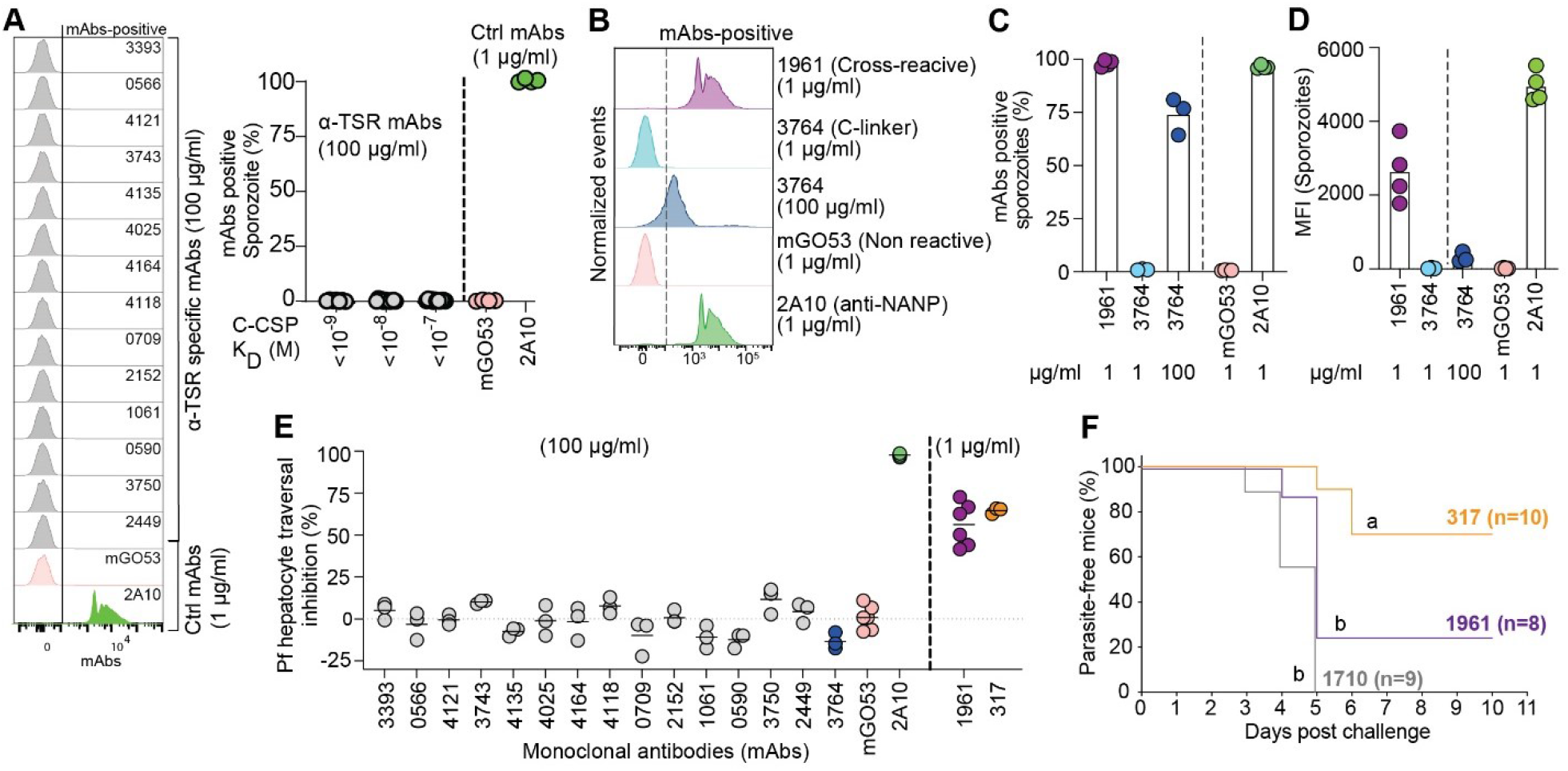
C-CSP-specific mAbs lack anti-parasite activity. **(A)** Flow-cytometric profiles (left) and percent of live (right) PbPfCSP(mCherry) sporozoites recognized by α-TSR-specific mAbs (100 µg/ml) with the indicated affinities as determined by flow cytometry. The positive control anti-NANP mAb 2A10 (1 µg/ml; (Triller *et al*, 2017)) and negative control non-PfCSP reactive mAb mGO53 (1µg/ml; (Wardemann *et al*, 2003)) are shown for comparison (n=2-3). **(B)** Representative flow cytometry profiles, **(C)** percent of mAb-positive live *Pb-PfCSP* sporozoites and **(D)** mean fluorescence intensity (MFI) for the junction cross-reactive mAb 1961, C-linker-specific mAb 3764 and the control mAbs 2A10 and mGO53 (A) at the indicated concentrations (n=3-4). **(E)** Pf sporozoite hepatocyte traversal inhibitory capacity of the indicated C-CSP reactive, C-CSP cross-reactive (1961) and control (2A10, mGO53 and anti-NANP 317 mAb (Oyen *et al*, 2017) mAbs at the indicated concentrations (n=3-6). **(F)** Capacity of the passively-transferred indicated antibodies (100 μg) to protect mice (n=10 for mAb 317, n=8 for mAb 1961, and n=9 for mAb 1710) from parasitemia after the bite of three PbPfCSP(mCherry)-infected mosquitoes compared to the C-CSP-specific mAb 1710 (Scally *et al*., 2018). Data show the percentage of parasite-free mice in two independent experiments. Groups marked with the same letter were not statistically significantly different (Mantel–Cox log-rank test, see also Figure S5). Circles in **A, C-E** indicate independent experiments.

Thus, regardless of their fine epitope conformational specificity and affinity, anti-C-CSP antibodies lacked measurable sporozoite-binding capacity and *in vitro* parasite-inhibitory activity even at 100-fold higher concentrations than anti-repeat and cross-reactive mAbs. Only a single C-CSP reactive mAb with cross-reactivity to the repeat and N-terminal junction mediated parasite inhibition *in vitro* but not *in vivo*.

### mAb 3764 binds a DPN core sequence in PfCSP_281-294_

The C-linker specific mAb 3764 that weakly bound sporozoites was the only C-CSP specific mAb (Fig. S5A) encoded by *IGHV3-33*, a gene which is associated with germline NANP reactivity and also encodes for antibodies with cross-reactivity to NPDP and NVDP motifs in the PfCSP N-terminal junction (Murugan *et al*., 2020). The DPN motif in the linear target peptide P2 of mAb 3764 (QGHNMPN**DPN**RNVD; Fig. 3B, Fig. S3B) shared some similarity with a motif in the N-terminal junction (NP**DPN**AN). To define the molecular basis for the recognition of the C-terminal linker region, we solved the crystal structure of the 3764 Fab in complex with P2 to 2.36 Å resolution (Table S7). Epitope recognition is mediated by all mAb 3764 Fab complementarity-determining regions (CDRs) with the exception of the Igκ CDR2 (KCDR2, Fig. 5A.) The core P2 motif, DPN, interacts primarily with the IgH chain CDRs 2 and 3 (HCDR2 and 3, Fig. 5A, inset I), forming two hydrogen bonds and contributing 274.7 Å^2^ of buried surface area (BSA). Comparison of this binding mode to the core motif with that of mAb 4498, a VH3-33 mAb that binds to the DPN motif in the N-terminal junction (peptide NANPNV<u>DPN</u>ANP; PDB ID: 6ULF; (Murugan *et al*., 2020) shows that both mAbs recognize the DPN core in a highly similar disposition (whole atom RMSD of 0.24 Å; Fig. S5B and S5C). In periphery of the core epitope motif, P2 residues Met 285 (M285), Asp 287 (N287) and Arg 291 (R291) partake in important interactions: M285 contributes 160.3 Å^2^ of BSA at the interface with HCDR2 and KCDR3 (Fig. 5A, inset II); N287 (Fig. 5A, inset III) is responsible for five hydrogen bonds with HCDR3, KCDR1 and KCDR3 and contributes 156.8 Å^2^ of BSA; while R291 (Fig. 5A, inset IV) contributes four hydrogen bonds with HCDR1 and 106.3 Å^2^ of BSA. Sequence analysis reveals that mAb 3764 had undergone limited somatic hypermutation, with amino acid changes from the germline sequence in the VH (four in HCDR2 and one in the framework region 3 (FWR3) and Vκ (one in KCDR1 and two in the FWR3) chain. Of these mutated residues, H.N56Y and H.Y58S in the HCDR2 form part of the peptide-binding interface, together contributing 70.8 Å^2^ of BSA around M285 (Fig. 5A, inset II). Given the sequence similarities between the core DPN regions of the C-CSP peptide and the N-junction peptide (Fig. 5B), we investigated the molecular basis for mAb 3764-binding specificity to C-CSP. Modeling was performed to mutate M285, N287 and R291 residues (see Fig. 5A) to their corresponding N-junction peptide residues, Gly, Pro and Ala, respectively (Fig. 5B). The replacement of M285 with Gly would result in a much smaller BSA in the mAb 3764 HCDR2 and KCDR3. Replacing N287 with Pro would result in the loss of four of the five hydrogen bonds with HCDR3, KCDR1 and KCDR3, as well as introducing steric hindrance with KCDR1 (Fig. 5B). Finally, Ala in place of Arg in position 291 would result in a loss of three of the four hydrogen bonds with HCDR1. Taken together, the crystal structure resolving PfCSP residues 284 – 293 in complex with mAb 3764, and molecular modeling, helped to explain the specificity of mAb 3764 for C-CSP through interactions with the IgH (PfCSP_284-292_) and Igκ chains (PfCSP_285-287_ and PfCSP_289_) and lack of cross-reactivity with the DPN motif in the N-terminal junction.

**Figure 5.**
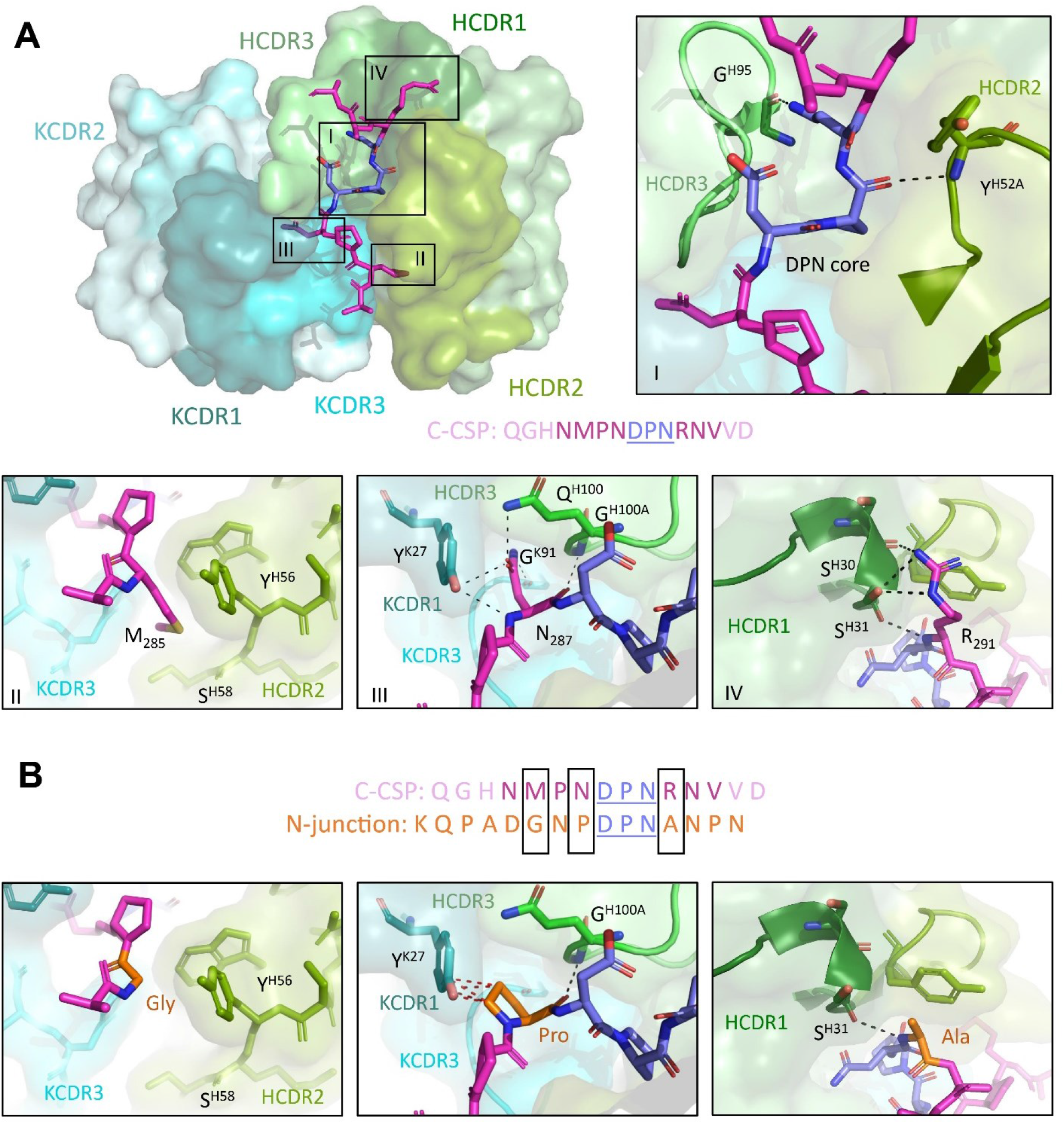
Molecular characterization of mAb 3764 recognition of the PfCSP. **(A)** Crystal structure showing the variable regions of the C-linker specific mAb 3764 and P2, with the mAb 3764 heavy chain illustrated in green and the kappa chain in blue. HCDR1, 2 and 3 (dark, pea and light green, respectively) as well as KCDR1 and 3 (dark teal and aqua, respectively) contribute to epitope recognition, but KCDR2 (teal) does not. The central DPN core (deep blue) of the C-CSP peptide (magenta) interacts with HCDR2 and 3 similarly to other VH3-33 antibodies (inset I), forming hydrogen bonds (H-bonds, shown with black dotted lines) with HCDR2 and 3. C-CSP residues M285 (inset II), N287 (inset III) and R291 (inset IV) contribute substantial buried surface area (BSA) to the complex. (II) M285 does not contribute any H-bonds, but does contribute to BSA. N287 (III) and R291 (IV) are responsible for hydrogen bonding with HCDR1 or HCDR3, KCDR1 and KCDR3 respectively. **(B)** Despite significant sequence similarity in the region of the DPN core between the C-CSP peptide and the N-junction peptide (sequence shown in orange), mAb 3764 does not bind to the N-junction peptide. Molecular modeling replacing M285, N287 and R291 with Gly, Pro and Ala, respectively, reveals these alternate residues would result in a loss of hydrogen bonds and reduced BSA.

## DISCUSSION

The characterization of our large panel of human mAbs provides insights in the gene usage characteristics, development, cellular origin, epitope specificity, binding mode and functional properties of C-CSP reactive antibodies. Similar to the strong association between NANP-repeat reactivity and *IGHV3-33/IGKV1-5* gene usage, (Imkeller *et al*., 2018; Murugan *et al*., 2018; Tan *et al*., 2018), our data establish a link between C-CSP-reactivity and *IGHV3-21* and *IGVL3-21 or IGVL3-1* gene usage for Th2R/Th3R specific antibodies, which might allow conclusions about the frequency of C-CSP-reactive cells based on Ig gene sequencing. The association between *IGHV3-21* gene usage and antibody specificity for the C-CSP α-TSR domain appears to be independent of whether the immune system responds to recombinant protein or whole sporozoites, since Ig genes encoding VH3-21 anti-C-CSP mAbs have also been cloned from B cells isolated from RTS,S/AS01 and PfSPZ CVac vaccinees (Beutler *et al*., 2022; Scally *et al*., 2018). Nevertheless, the high *IGHV3-21* frequency that we observed after immunization with 3 doses of the PfSPZ Vaccine (900,000 radiation attenuated PfSPZ administered on days 1, 8, and 29) compared to immunization with the chemo-attenuated PfSPZ-CVac (51,200 PfSPZ administered on days 1, 29, and 57 under chloroquine; Mordmuller *et al*, 2017) with strong dominance of *IGHV3-33* (Imkeller *et al*., 2018; Murugan *et al*., 2018; Tan *et al*., 2018) suggests that radiation, dose, or dosing interval might influence the anti-sporozoite B cell response. The large number of VH3-21 and non-VH3-21 B cells with Th2R/Th3R specificity including many that expressed unmutated antibodies indicates that the naïve B cell repertoire contains potent anti-Th2R/Th3R precursors. The relative abundance of these cells might contribute to the apparent immunodominance of this epitope compared to the conserved RII+, which is targeted by antibodies with diverse gene combinations, and compared to non-α-TSR epitopes.

In contrast to antibodies against the α-TSR domain, antibodies with reactivity to the C-linker appear to be rare and target linear epitopes. The differences in abundance of these antibodies may be linked to differences in the frequency of naïve precursor B cells that recognize these epitopes and structural differences between the C-linker region and α-TSR domain or differences in epitope presentation and immunogenicity. Analysis of the crystal structure of mAb 3764 in complex with amino acid residues 284-293, provides first insights into how the C-linker domain is recognized by a non-cross-reactive human anti-PfCSP antibody. Despite sequence identity of the core DPN epitope in the C-linker with the N-terminal junction, the structure data provide a molecular explanation for the specificity of mAb 3764 for C-CSP. Steric constraints preclude cross-reactivity with the junction in contrast to mAb 1961, which shows flexibility in binding to peptides in the C-linker, repeat and junction, likely due to recognition of the similar 4-aa NANA, NANP, NPDP and NVDP motifs in these domains. The data support the notion that antibodies recognize a core motif, but that the context of their complete epitope including peripheral residues defines their binding specificity and cross-reactivity.

Despite the relative abundance of germline-encoded α-TSR-reactive antibodies, B cells with this specificity seem to undergo efficient affinity maturation and IgG isotype switching in germinal center reactions. The relative abundance of IgG compared to IgM antibodies and higher numbers of selected somatic mutations in VH3-21 mAbs compared to VH3-33 mAbs, and enrichment of affinity-increasing H.T33 mutation, suggests that epitope specificity is linked to differences in cell differentiation pathways or cell differentiation kinetics. The large number of NANP motifs in the central repeat domain might mediate stronger BCR cross-linking and B cell activation signals compared to the non-repeating epitopes in the PfCSP C terminus, thereby affecting B cell activation, differentiation and germinal center selection (Imkeller *et al*., 2018). Alternatively, and not mutually exclusively, structure rather than valency differences between the disordered and highly flexible central repeat region compared to the structured C-CSP domain may underlie these isotype and mutation-load differences (MacRaild *et al*, 2018). Future studies will have to define the molecular and cellular mechanisms and determine whether epitope-associated differences in affinity maturation between repeat and C-CSP antibodies are also seen in response to immunization with recombinant PfCSP and RTS,S/AS01 or other antigens with repeating disordered domains that are characteristic of numerous parasite proteins.

Our study shows that the C-CSP of irradiated sporozoites is highly immunogenic and identifies linear target epitopes in the C-terminal linker and α-TSR domain. However, in assays with live Pf sporozoites none of the C-CSP epitopes seem to be the target of potent parasite-inhibitory antibodies. Even the C-linker specific mAb 3764, which showed weak parasite binding, lacked Pf-inhibitory activity at a hundred-fold higher concentration than antibodies against the central repeat domain, suggesting that modest C-CSP binding is not sufficient to mediate protection. Although the C-linker region may be slightly easier to reach for C-CSP mAbs than the α-TSR, which is directly anchored to the parasite surface, the central repeat and junction domains are clearly more accessible on the surface of live sporozoites. The direct comparison of C-CSP and NANP antibodies demonstrates that the almost complete inaccessibility of the PfCSP C-terminal domain strongly limits the direct parasite-inhibitory activity and potency of antibodies with specificity for this domain.

We have shown previously that passive transfer of the Th2R/Th3R-reactive mAb 1710 at high dose fails to protect mice from infection with PfCSP transgenic *P. berghei* parasites and the development of blood stage parasitemia (Murugan *et al*., 2020; Scally *et al*., 2018), findings that have recently been confirmed in an independent study (Wang *et al*., 2021). Since the epitopes of the Th2R/Th3R antibodies identified in this study overlap with the mAb 1710 target site (including many that share VH3-21/Vλ3-21 Ig gene combinations), we do not expect these mAbs to exhibit any direct parasite-inhibitory activity *in vivo*. Given their lack of Pf sporozoite binding and *in vitro* parasite inhibition, we assume that the RII+ reactive antibodies would behave similarly. Some degree of *in vivo* protection from i.v. parasite challenge has recently been reported after high-dose passive transfer of the RII+ and the Th2R/Th3R-reactive mAbs 1512 and 236, respectively (Beutler *et al*., 2022). How the potency of these high-affinity mAbs compares to anti-repeat mAbs with similar binding strength has not been determined but given that mAbs 1512 and 236 target the same epitopes as the mAbs in our collection, we expect it to be overall low.

In response to RTS,S/AS01 immunization, anti-NANP and anti-C-terminus antibody titers correlate with protection (Chaudhury *et al*, 2021; Chaudhury *et al*, 2016; Dobano *et al*, 2019). To what degree the C-CSP-reactive serum antibodies recognize C-CSP specifically or cross-react with the repeat and junction has not been determined. Therefore, it is unclear whether protection is associated with C-CSP-specific or cross-reactive antibodies. Passive transfer of mAb mAb1961 shows that cross-reactive mAbs can mediate low levels of parasite inhibition *in vivo*. Such antibodies likely confer some degree of protection by binding to repeat and junction epitopes that are readily accessible on the surface of live sporozoites and not to the hidden C-CSP (Murugan *et al*., 2020). Cross-reactive anti-PfCSP mAbs have been shown to have overall higher affinity than antibodies that target the junction, repeat or C-CSP specifically (Murugan *et al*., 2020). Thus, protection seems to be linked to high affinity C-CSP cross-reactive rather than C-CSP specific antibodies. Nevertheless, we cannot exclude that C-CSP specific antibodies gain access to the C-terminus upon PfCSP binding by high affinity anti-NANP or junction antibodies. PfCSP shedding induced by anti-repeat/junction antibodies might uncover epitopes in the α-TSR domain enabling C-CSP-antibody binding and parasite clearance. Although synergistic effects of anti-repeat/junction and anti-C-CSP mAbs may not be easily detected in animal models with human monoclonal antibodies (Wang *et al*., 2021), antibody effector functions such as opsonization and phagocytosis or complement activation may contribute to parasite inhibition in humans. However, the high sequence diversity of the Th2R/Th3R domain among Pf parasite populations in natural settings will limit these indirect protective effects to antibodies against the conserved RII+ of the α-TSR domain, and to protection from parasites with high sequence similarity to the vaccine strain.

In summary, with our mAb collection we demonstrate the lack of accessible C-CSP epitopes on live sporozoites as targets for potent parasite-inhibitory antibodies. Due to the high immunogenicity of the PfCSP C terminus, these observations are of direct relevance for the design of an improved PfCSP-based malaria vaccine.

## MATERIALS and METHODS

### Clinical samples

Serum and peripheral blood mononuclear cells (PBMCs, obtained by Ficoll density gradient centrifugation) were collected from malaria-naïve, healthy European volunteers who participated in the MAVACHE clinical verification study with three doses of 900,000 PfSPZ four weeks apart (Mordmuller *et al*., 2022). Ethical approval was obtained from the ethics committee of the medical faculty and the university clinics of the University of Tübingen. The study was conducted according to the principles of the Declaration of Helsinki.

### Cell lines

Human embryonic kidney HEK293T cells (Invitrogen, 0.3 × 10^6^ /ml) were cultured under rotating conditions (180 rpm) in 50ml bioreactors with FreeStyle 293-F medium at 37°C and 8% CO_2_. HC-04 cells (Sattabongkot *et al*, 2006) were cultured at 37°C and 5% CO_2_ in HC-04 complete medium with 428.75 ml MEM (without L-glutamine), 428.75 ml F-12 Nutrient Mix (with L-glutamine), 15 mM HEPES, 1.5 g/l NaHCO3, 2.5 mM L-glutamine, 10% FCS and 1% penicillin/streptomycin solution.

### Bacteria

MAX efficiency DH10B™ competent cells were grown in LB medium for cultivation and in terrific broth for plasmid purification. Bacterial shaker at 37 °C and 180 rpm was used for cultivation.

### Proteins and peptides

Biotinylated NF54 FL-CSP, non-biotinylated 7G8 FL-CSP and α-TSR protein were expressed in HEK293F cells and purified by HisTrap HP (Cytiva) and size exclusion chromatography (Superdex 200 Increase 10/300 GL, Cytiva). FL-CSP and C-CSP were expressed in *E*.*coli* by the EMBL protein expression and purification core facility (Heidelberg, Germany). N-junc, NANP_5_, NANP_10_, C-linker and the 14 overlapping peptides (P1-14) covering the complete NF54 PfCSP C-terminus were synthesized by PSL Peptide Specialty Laboratories GmbH (Heidelberg, Germany). Sequences of peptides and proteins are listed in Table S1.

### Parasites, mosquitoes and mice

*Anopheles coluzzii* (Ngousso strain) and transgenic *Anopheles gambiae* (*7b* strain (Pompon & Levashina, 2015)) were maintained at 28 °C 70-80% humidity 12/12 h day/night cycle. The *P. falciparum* NF54 clone (*Pf*) used in this study originated from Prof. Sauerwein’s laboratory (RUMC, Nijmegen) and was regularly tested for *Mycoplasma* contamination. For *P. falciparum* infections, *A. coluzzii* mosquitoes were fed for 15 min via artificial midi feeders (Glass Instruments, The Netherlands) with NF54 gametocyte cultures and kept at 26°C 80% humidity 12/12 h day/night cycle in a secured S3 laboratory according to the national regulations (Landesamt für Gesundheit und Soziales, project number 297/13). Infected mosquitoes were offered an additional uninfected blood meal to boost sporozoite formation eight days post infection (dpi). Sporozoites were dissected for traversal assay 5-7 days later as described below.

*A. gambiae 7b* mosquitoes, an immunocompromised transgenic mosquito line derived from the G3 laboratory strain, were used for the production of transgenic *P. berghei* sporozoites expressing PfCSP and the reporter fluorescence protein mCherry (PbPfCSP(mCherry)). Briefly, female CD1 mice (7-12-weeks-old) and female C57BL/6J mice (8-weeks-old) were bred in the MPIIB Experimental Animal Facility (Marienfelde, Berlin) and housed in a pathogen-free animal facility in accordance with the German Animal Protection Law (§8 Tierschutzgesetz) and approved by the Landesamt für Gesundheit und Soziales (LAGeSo), Berlin, Germany (project numbers 368/12 and H0335/17). *A. gambiae 7b* mosquitoes were fed on PbPfCSP(mCherry)-infected female CD1 mice 3 days post passage (0.1-0.8% gametocytemia) for 30-45 min, and kept at 20°C 80% humidity 12/12 h day/night cycle. Infected mosquitoes were offered an additional uninfected blood meal to boost sporozoite formation 7 dpi, and after eleven days were used for C57BL/6J mice challenge experiments and live sporozoite FACS staining as described below.

### Flow cytometry and isolation of PfCSP-specific memory B cells

Frozen PBMCs were thawed, washed with RPMI 1640 (Gibco) and incubated with full-length biotinylated NF54 PfCSP in the presence of the following mouse anti-human antibodies at the indicated dilutions: CD27-phycoerythrin (PE) (M-T271) at 1:5, CD38-Brilliant Violet 605 (BV605) (HB7) at 1:20, IgG-BV510 (G18-145) at 1:20, IgM-BV421 (G20-127) at 1:20, IgD-Allophycocyanin-H7 (APC-H7) (IA6-2) at 1:20 (all from BD biosciences), CD19-(BV786) (HIB19) at 1:10 and CD21-PE-Cy7 (Bu32) at 1:20 (both from biolegend). Biotin was detected using streptavidin FITC (1:1000). 7-Aminoactinomycin D (7-AAD) (Invitrogen) at 1:400 was used as a marker for dead cells. Single-cell sorting of 7AAD−PfCSP+CD19+ B cells gated positive for either IgG or CD27 as PfCSP memory B cells and 7AAD−CD38+CD27+CD19+ cells as plasmablasts was performed using a FACS Aria II (BD Biosciences) with FACSDiva software (version 8.0.1) using the index sort option. Data were analyzed using FlowJo v.10.0.8 (Tree Star).

### Ig gene amplification and sequence analysis

A high-throughput robotic platform was used to amplify the *IGH, IGK* and *IGL* loci of Ig genes by RT-PCR as previously reported (Busse *et al*, 2014; Murugan *et al*, 2015; Tiller *et al*, 2008). Briefly, after cDNA synthesis with random hexamers, a nested polymerase chain reaction (PCR) with barcoded primers in the second PCR was used to amplify the heavy and light chain genes. Amplicons were pooled, purified, and sequenced using the MiSeq 2 × 300 base pair (bp) paired-end sequencing platform (Illumina). Integrated indexed flow cytometry and sequence data integration as well as Ig gene annotation was performed using sciReptor version 1.0-6-g034c8ae (Busse *et al*., 2014; Imkeller *et al*, 2016). Downstream sequence analyses and data visualization were performed using GraphPad Prism (version 9.12) and R 4.2.2 and ggplot2.

### Cloning of Ig genes and recombinant monoclonal antibodies production

Cloning of the Ig genes and recombinant antibody production were performed as previously described (Tiller *et al*, 2008). Ig genes were amplified with primers containing restriction site-tagged V- and J-specific. The amplicons were cloned into human *IGHG1* or *IGKC* or *IGLC* expression vectors (Addgene number 80795, 80796 and 99575, respectively). After co-transfection of the heavy and light expressing plasmid DNA into HEK293T cells (Invitrogen), the recombinant monoclonal antibodies were purified from the supernatant with protein G beads (GE Healthcare).

### Enzyme Linked Immunosorbent assay (ELISA)

Serum, antigen and polyreactivity ELISAs were performed as described (Murugan *et al*., 2020; Triller *et al*., 2017). In brief, 384-well polystyrene plates (Corning) with high binding capacity were coated overnight (O/N) at 4 °C with full-length PfCSP (0.4 µg/ml), N-junc, NANP_5_, NANP_10_ (2 µg/ml), C-CSP (1 µg/ml), Insulin (10 ug/ml; Sigma Aldrich), lipopolysaccharide (LPS; 10 ug/ml; Sigma Aldrich) and double-stranded DNA (dSDNA; 20 ug/ml; Sigma Aldrich) in PBS. The plates were washed 3 times with PBS-T (0.05% Tween20 in PBS) using a Tecan plate washer. ELISA plates were blocked at RT for 1 hour with 4% BSA in PBS (serum ELISA), 1% BSA in PBS (antigen ELISA) or blocking buffer (1x PBS, 0.05% (v/v) Tween20, 1 mM EDTA, for polyreactivity ELISA). Serum samples serially diluted at an initial dilution of 1:200 in 1% BSA with PBS, or mAbs at an initial concentration of 4 µg/ml (antigen ELISA) or 1 µg/ml (polyreactivity ELISA) in 1x PBS were loaded on the plate and incubated at RT for 1.5 hours. Antibodies were detected with goat anti-human IgG secondary antibody coupled to horseradish peroxidase (HRP) (Jackson Immuno Research) diluted at 1:1,000 in blocking buffer (1x PBS, 0.05% (v/v) Tween20, 1 mM EDTA) and then detected with 2,2’-azino-bis-(3-ethylbenzothiazoline-6-sulfonic acid) diammonium (ABTS) substrate (Roche Diagnostics) diluted at 1:1000 in H_2_O_2_. Optical density (OD) at 405 nm was determined using an M1000Pro plate reader (Tecan). Area under the curve (AUC) values were calculated using GraphPad Prism 9.1.2. Serum from placebo recipients was used as a negative control in serum ELISAs. mAbs 2A10 (Triller *et al*., 2017) and mGO53 (Wardemann *et al*, 2003) were used as positive and negative controls, respectively, in antigen ELISAs and mAbs ED38 (Meffre *et al*, 2004) and mGO53 as positive and negative controls, respectively, in poly-reactivity ELISAs.

### Blocking ELISA

Blocking ELISA was performed as described above for antigen ELISA, with the following modifications. After 1.5 hours incubation with the mAbs (starting concentration of 64 μg/ml in 1X PBS and 1:2 dilution) at room temperature, biotinylated mAb 1710 (Scally *et al*., 2018) or mAb 1512 (Beutler *et al*., 2022) at 0.5 μg/ml was added to compete for an additional 15 mins at room temperature. mAb 1710 or mAb 1512 binding was detected with streptavidin-HRP diluted at 1:1,000 in blocking buffer (1x PBS, 0.05% (v/v) Tween20, 1 mM EDTA) and then detected with 2,2’-azino-bis-(3-ethylbenzothiazoline-6-sulfonic acid) diammonium (ABTS) substrate (Roche Diagnostics) diluted at 1:1000 in H_2_O_2_. The optimal concentrations of the biotinylated 1710 and 1512 were determined by measuring their ability to self-block their non-biotinylated versions over a range of concentrations.

### Surface Plasmon Resonance (SPR)

SPR measurements were performed using a Biacore T200 (GE Health-care) instrument docked with an S CM5 series sensor chip (GE Healthcare), as described (Murugan *et al*., 2018). In brief, anti-human IgG antibodies were immobilized on the chip using an amine coupling-based human antibody Fab capture kit. Hepes (10 mM) with 150 mM NaCl at pH 7.4 was used as running buffer. Equal concentrations of the sample antibody and isotype control (mGO53 (Wardemann *et al*., 2003)) were captured in the sample and reference flow cells, respectively. Running buffer was injected at a rate of 10 μl/min for 20 min to stabilize the flow cells. C-CSP peptide at 0.015, 0.09, 0.55, 3.3 and 20 μM in running buffer was injected at a rate of 30 μl/min. The flow cells were regenerated with 3 M MgCl_2_. Data were fitted by steady-state kinetic analysis using BIACORE T200 software V2.0.

### mAb 3764 Fab production

mAb 3764 heavy and light chain variable regions were cloned into custom pcDNA3.4 expression vectors upstream of human IgK and IgG1-C_H_1 domains, and transiently co-transfected into HEK293F cells. The resulting Fab was purified by Kappa-select affinity chromatography (Cytiva), cation exchange chromatography (MonoS; Cytiva), and size exclusion chromatography (Superdex 200 Increase 10/300 GL; Cytiva).

### Crystallization and structure determination

Purified 3764 Fab and PfCSP C-terminal linker peptide (PfCSP_281-294_) were mixed in a 1:3 molar ratio; and, in sparse matrix screening, crystals of the complex emerged from a 1:1 ratio mix with 19% (v/v) isopropanol, 19% (w/v) PEG 4000, 5% (v/v) glycerol, 0.095 M sodium citrate, pH 5.6. Crystals did not require cryopreservation before being flash frozen in liquid nitrogen. Data was collected at the 23-ID-B beamline at the Advanced Photon Source and manually processed using XDS (Kabsch, 2010). Molecular replacement using the heavy and light chains from the 4498 Fab (PDB ID: 6O24) was carried out in Phaser (McCoy *et al*, 2007). Iterative refinement was performed in Coot (Emsley *et al*, 2010) and phenix.refine (Adams *et al*, 2010). All software were accessed through SBGrid (Morin *et al*, 2013). The structure has been deposited in the PDB and is accessible under ID: 8FG0.

### Pf sporozoite hepatocyte traversal assay *in vitro*

Traversal assays were performed as previously described (Murugan *et al*., 2018; Triller *et al*., 2017). Briefly, Pf sporozoites were isolated from the mosquito salivary glands 13-15 days post-infection in complete HC-04 medium, pre-incubated with monoclonal antibody at the indicated concentrations for 30 min on ice and added to the human hepatocyte cell line (HC-04 (Sattabongkot *et al*., 2006)) for 2 h at 37 °C and 5% CO2 in the presence of 0.5 mg/ml dextran-rhodamine (Molecular Probes). Untreated Pf sporozoites were used to measure maximum traversal capacity. A chimeric version of mAb 2A10 (Zavala *et al*., 1983) with human IgG1 constant region (Triller *et al*., 2017), and mGO53 (Wardemann *et al*., 2003) were used as positive and negative controls, respectively. Cells were trypsinized, washed and fixed with 1% PFA in PBS before measuring dextran positivity using the FACS LSR II instrument (BD Biosciences). Data analysis was performed by subtracting background (dextran positivity of cells treated with uninfected mosquito salivary gland material) and normalizing to maximum Pf traversal capacity (dextran positivity of cells without mAbs) using FlowJo V.10.0.8 (Tree Star).

### *Pb-PfCSP*(mCherry) live sporozoite flow cytometry

*Plasmodium berghei* PbPfCSP(mCherry) sporozoites were isolated from mosquito salivary glands 18 days post infection in HC-04 complete medium, incubated for 30 min on ice in siliconized tubes with the indicated concentrations of mAbs (150,000 sporozoites in total volume of 100 μl 1% FCS/PBS). Sporozoites were washed with 1.5 ml of 1% FCS/PBS by centrifugation at 9,300g for 4 min at 4°C. The sporozoite pellet was resuspended in 100 μl 1% FCS/PBS containing anti-human IgG1-Cy5 (2 µg/ml, Central Laboratory Facility Deutsches Rheuma-Forschungszentrum, Berlin) and incubated on ice for 30 min. The sporozoite suspension was washed again, and the pellet was re-suspended in 200 μl 1% FCS/PBS. Sporozoites stained with antibodies were detected by endogenous mCherry expression and analyzed for Cy5 signal in a FACS LSR II instrument (Becton Dickinson). Salivary gland material from uninfected mosquitoes was used as sporozoite gating control (Fig. S4A). Data were analyzed using FlowJo V.10.0.8 (Tree Star).

### *P. berghei* PbPfCSP(mCherry) mouse challenge model by mosquito bites

Female C57BL/6J mice were passively immunized by intraperitoneal injection of 100 μg of monoclonal antibodies in 100-200 μL PBS. After 20 h, mice were exposed to the bites of three PbPfCSP(mCherry) salivary gland-infected mosquitoes as described elsewhere (Ludwig et al. in preparation). Briefly, PbPfCS*P*(mCherry)-infected mosquitoes were put to sleep on ice at 17 dpi and selected for mCherry signal in the salivary glands under a fluorescence stereo microscope (Leica, M205 FA). Single salivary gland-positive mosquitoes were transferred into individual containers and starved overnight. The next day (18 dpi), anesthetized naïve C57BL/6J mice were exposed for 10 min to three bites of the selected PbPfCSP(mCherry)-infected mosquitoes. Peripheral whole blood was collected from the submandibular vein 2-3 h after challenge to isolate sera to quantify concentration mAbs titers by ELISA. From days 3 to 7 and on day 10 post mosquito bite, parasitemia (mCherry-positive red blood cells / total red blood cells) was measured by flow cytometry (LSR II instrument, BD Biosciences), and confirmed by Giemsa-stained thin blood smears. All infected mice were euthanized on day 7 post mosquito bite, before the occurrence of malaria symptoms. FACS data were analyzed by FlowJo V.10.0.8 and the pre-patency period was declared on the first day when parasitemia values were above the background signal of negative mice.

### Statistics

Statistical analysis was performed using GraphPad Prism (version 9.12). The corresponding statistical test used for each experiment is stated in the figure legend (\**P* < 0.05, \*\**P* < 0.01, \*\*\**P* < 0.001, and \*\*\*\**P* < 0.0001).

## ACKNOWLEDGEMENTS

The authors thank Dorien Foster, Julia Gärtner, Christine Niesik, Claudia Winter (DKFZ) and H. Gordon, C. Kreschel, M. Andres, D. Eyermann and L. Spohr (Max Planck Institute for Infection Biology, Berlin) for experimental assistance, Christian E. Busse (DKFZ) for bioinformatic support, the EMBL Protein Expression and Purification Core Facility and the DKFZ/ EMBL/Heidelberg University Chemical Biology Core Facility, especially P. Sehr, for technical assistance and services. The work was supported by the Bill and Melinda Gates Foundation (OPP1179906; J.-P.J, H.W. and E.A.L.), by the CIFAR Azrieli Global Scholar program (J.P.J.), the Ontario Early Researcher Award program (J.P.J.), and the Canada Research Chair program (J.P.J.). C.B.A was supported by a by Hospital for Sick Children Restracomp Postdoctoral Fellowship and a Banting Postdoctoral Fellowship. X-ray diffraction experiments were performed at GM/CA@APS, which has been funded in whole or in part with federal funds from the National Cancer Institute (ACB-12002) and the National Institute of General Medical Sciences (AGM-12006). The Eiger 16 M detector at GM/CA-XSD was funded by NIH grant S10 OD012289. This research used resources of the Advanced Photon Source, a US Department of Energy (DOE) Office of Science user facility operated for the DOE Office of Science by Argonne National Laboratory under contract DE-AC02-06CH11357. Manufacture of PfSPZ Vaccine was funded in part by the National Institute of Allergy and Infectious Diseases of the National Institutes of Health under SBIR award numbers 5R44AI058375 and 5R44AI055229.

## AUTHORS CONTRIBUTION

O.E.O., G.C., and C.B.A. designed, conducted the experiments and interpreted experimental results. O.E.O. and A.O. performed analyses and interpreted results. K.P. and C.C. conducted experiments. S.L.H., P.G.K., and B.M. produced the vaccine and/or provided clinical trial samples. R.M designed experiments and interpreted experimental results. H.W., E.A.L. and J.-P.J. conceived the study and interpreted experimental results. O.E.O., G.C., C.B.A., J.-P.J., E.A.L., and H.W. wrote the manuscript.

## REFERENCES

Adams PD, Afonine PV, Bunkoczi G, Chen VB, Davis IW, Echols N, Headd JJ, Hung LW, Kapral GJ, Grosse-Kunstleve RW et al (2010) PHENIX: a comprehensive Python-based system for macromolecular structure solution. Acta Crystallogr D Biol Crystallogr 66: 213–221

Agnandji ST, Lell B, Soulanoudjingar SS, Fernandes JF, Abossolo BP, Conzelmann C, Methogo BG, Doucka Y, Flamen A, Mordmuller B et al (2011) First results of phase 3 trial of RTS,S/AS01 malaria vaccine in African children. N Engl J Med 365: 1863–1875

Aragam NR, Thayer KM, Nge N, Hoffman I, Martinson F, Kamwendo D, Lin FC, Sutherland C, Bailey JA, Juliano JJ (2013) Diversity of T cell epitopes in Plasmodium falciparum circumsporozoite protein likely due to protein-protein interactions. PLoS One 8: e62427

Asante KP, Abdulla S, Agnandji S, Lyimo J, Vekemans J, Soulanoudjingar S, Owusu R, Shomari M, Leach A, Jongert E et al (2011) Safety and efficacy of the RTS,S/AS01E candidate malaria vaccine given with expanded-programme-on-immunisation vaccines: 19 month follow-up of a randomised, open-label, phase 2 trial. Lancet Infect Dis 11: 741–749

Beutler N, Pholcharee T, Oyen D, Flores-Garcia Y, MacGill RS, Garcia E, Calla J, Parren M, Yang L, Volkmuth W et al (2022) A novel CSP C-terminal epitope targeted by an antibody with protective activity against Plasmodium falciparum. PLoS Pathog 18: e1010409

Bojang KA, Milligan PJ, Pinder M, Vigneron L, Alloueche A, Kester KE, Ballou WR, Conway DJ, Reece WH, Gothard P et al (2001) Efficacy of RTS,S/AS02 malaria vaccine against Plasmodium falciparum infection in semi-immune adult men in The Gambia: a randomised trial. Lancet 358: 1927–1934

Casares S, Brumeanu TD, Richie TL (2010) The RTS,S malaria vaccine. Vaccine 28: 4880–4894

Cawlfield A, Genito CJ, Beck Z, Bergmann-Leitner ES, Bitzer AA, Soto K, Zou X, Hadiwidjojo SH, Gerbasi RV, Mullins AB et al (2019) Safety, toxicity and immunogenicity of a malaria vaccine based on the circumsporozoite protein (FMP013) with the adjuvant army liposome formulation containing QS21 (ALFQ). Vaccine 37: 3793–3803

Chaudhury S, MacGill RS, Early AM, Bolton JS, King CR, Locke E, Pierson T, Wirth DF, Neafsey DE, Bergmann-Leitner ES (2021) Breadth of humoral immune responses to the C-terminus of the circumsporozoite protein is associated with protective efficacy induced by the RTS,S malaria vaccine. Vaccine 39: 968–975

Chaudhury S, Ockenhouse CF, Regules JA, Dutta S, Wallqvist A, Jongert E, Waters NC, Lemiale F, Bergmann-Leitner E (2016) The biological function of antibodies induced by the RTS,S/AS01 malaria vaccine candidate is determined by their fine specificity. Malar J 15: 301

Coppi A, Pinzon-Ortiz C, Hutter C, Sinnis P (2005) The Plasmodium circumsporozoite protein is proteolytically processed during cell invasion. J Exp Med 201: 27–33

Dame JB, Williams JL, McCutchan TF, Weber JL, Wirtz RA, Hockmeyer WT, Maloy WL, Haynes JD, Schneider I, Roberts D et al (1984) Structure of the gene encoding the immunodominant surface antigen on the sporozoite of the human malaria parasite Plasmodium falciparum. Science 225: 593–599

Dobano C, Sanz H, Sorgho H, Dosoo D, Mpina M, Ubillos I, Aguilar R, Ford T, Diez-Padrisa N, Williams NA et al (2019) Concentration and avidity of antibodies to different circumsporozoite epitopes correlate with RTS,S/AS01E malaria vaccine efficacy. Nat Commun 10, 2174 (2019) 10: 2174

Doud MB, Koksal AC, Mi LZ, Song G, Lu C, Springer TA (2012) Unexpected fold in the circumsporozoite protein target of malaria vaccines. Proc Natl Acad Sci U S A 109: 7817–7822

Emsley P, Lohkamp B, Scott WG, Cowtan K (2010) Features and development of Coot. Acta Crystallogr D Biol Crystallogr 66: 486–501

Espinosa DA, Gutierrez GM, Rojas-Lopez M, Noe AR, Shi L, Tse SW, Sinnis P, Zavala F (2015) Proteolytic Cleavage of the Plasmodium falciparum Circumsporozoite Protein Is a Target of Protective Antibodies. J Infect Dis 212: 1111–1119

Gandhi K, Thera MA, Coulibaly D, Traore K, Guindo AB, Doumbo OK, Takala-Harrison S, Plowe CV (2012) Next generation sequencing to detect variation in the Plasmodium falciparum circumsporozoite protein. Am J Trop Med Hyg 86: 775–781

Genito CJ, Beck Z, Phares TW, Kalle F, Limbach KJ, Stefaniak ME, Patterson NB, Bergmann-Leitner ES, Waters NC, Matyas GR et al (2017) Liposomes containing monophosphoryl lipid A and QS-21 serve as an effective adjuvant for soluble circumsporozoite protein malaria vaccine FMP013. Vaccine 35: 3865–3874

Gordon DM, McGovern TW, Krzych U, Cohen JC, Schneider I, LaChance R, Heppner DG, Yuan G, Hollingdale M, Slaoui M et al (1995) Safety, immunogenicity, and efficacy of a recombinantly produced Plasmodium falciparum circumsporozoite protein-hepatitis B surface antigen subunit vaccine. J Infect Dis 171: 1576–1585

Gupta NT, Vander Heiden JA, Uduman M, Gadala-Maria D, Yaari G, Kleinstein SH (2015) Change-O: a toolkit for analyzing large-scale B cell immunoglobulin repertoire sequencing data. Bioinformatics 31: 3356–3358

Herrera R, Anderson C, Kumar K, Molina-Cruz A, Nguyen V, Burkhardt M, Reiter K, Shimp R, Jr., Howard RF, Srinivasan P et al (2015) Reversible Conformational Change in the Plasmodium falciparum Circumsporozoite Protein Masks Its Adhesion Domains. Infect Immun 83: 3771–3780

Hutter JN, Robben PM, Lee C, Hamer M, Moon JE, Merino K, Zhu L, Galli H, Quinn X, Brown DR et al (2022) First-in-human assessment of safety and immunogenicity of low and high doses of Plasmodium falciparum malaria protein 013 (FMP013) administered intramuscularly with ALFQ adjuvant in healthy malaria-naive adults. Vaccine 40: 5781–5790

Imkeller K, Scally SW, Bosch A, Marti GP, Costa G, Triller G, Murugan R, Renna V, Jumaa H, Kremsner PG et al (2018) Antihomotypic affinity maturation improves human B cell responses against a repetitive epitope. Science 360: 1358–1362

Kabsch W (2010) Xds. Acta Crystallogr D Biol Crystallogr 66: 125–132

Kisalu NK, Idris AH, Weidle C, Flores-Garcia Y, Flynn BJ, Sack BK, Murphy S, Schon A, Freire E, Francica JR et al (2018) A human monoclonal antibody prevents malaria infection by targeting a new site of vulnerability on the parasite. Nat Med 24: 408–416

MacRaild CA, Seow J, Das SC, Norton RS (2018) Disordered epitopes as peptide vaccines. Pept Sci (Hoboken) 110: e24067

McCoy AJ, Grosse-Kunstleve RW, Adams PD, Winn MD, Storoni LC, Read RJ (2007) Phaser crystallographic software. J Appl Crystallogr 40: 658–674

McCutchan TF, Kissinger JC, Touray MG, Rogers MJ, Li J, Sullivan M, Braga EM, Krettli AU, Miller LH (1996) Comparison of circumsporozoite proteins from avian and mammalian malarias: biological and phylogenetic implications. Proc Natl Acad Sci U S A 93: 11889–11894

Meffre E, Schaefer A, Wardemann H, Wilson P, Davis E, Nussenzweig MC (2004) Surrogate light chain expressing human peripheral B cells produce self-reactive antibodies. J Exp Med 199: 145–150

Mordmuller B, Sulyok Z, Sulyok M, Molnar Z, Lalremruata A, Calle CL, Bayon PG, Esen M, Gmeiner M, Held J et al (2022) A PfSPZ vaccine immunization regimen equally protective against homologous and heterologous controlled human malaria infection. NPJ Vaccines 2022 Aug 23;7(1):100 doi: 101038/s41541-022-00510-z 7: 100

Mordmuller B, Surat G, Lagler H, Chakravarty S, Ishizuka AS, Lalremruata A, Gmeiner M, Campo JJ, Esen M, Ruben AJ et al (2017) Sterile protection against human malaria by chemoattenuated PfSPZ vaccine. Nature 542: 445–449

Morin A, Eisenbraun B, Key J, Sanschagrin PC, Timony MA, Ottaviano M, Sliz P (2013) Collaboration gets the most out of software. Elife 2: e01456

Murugan R, Buchauer L, Triller G, Kreschel C, Costa G, Pidelaserra Marti G, Imkeller K, Busse CE, Chakravarty S, Sim BKL et al (2018) Clonal selection drives protective memory B cell responses in controlled human malaria infection. Sci Immunol 3, eaap8029 (2018)

Murugan R, Scally SW, Costa G, Mustafa G, Thai E, Decker T, Bosch A, Prieto K, Levashina EA, Julien JP et al (2020) Evolution of protective human antibodies against Plasmodium falciparum circumsporozoite protein repeat motifs. Nat Med 26: 1135–1145

Nardin EH, Nussenzweig V, Nussenzweig RS, Collins WE, Harinasuta KT, Tapchaisri P, Chomcharn Y (1982) Circumsporozoite proteins of human malaria parasites Plasmodium falciparum and Plasmodium vivax. J Exp Med 156: 20–30

Olotu A, Fegan G, Wambua J, Nyangweso G, Leach A, Lievens M, Kaslow DC, Njuguna P, Marsh K, Bejon P (2016) Seven-Year Efficacy of RTS,S/AS01 Malaria Vaccine among Young African Children. N Engl J Med 374: 2519–2529

Olotu A, Lusingu J, Leach A, Lievens M, Vekemans J, Msham S, Lang T, Gould J, Dubois MC, Jongert E et al (2011) Efficacy of RTS,S/AS01E malaria vaccine and exploratory analysis on anti-circumsporozoite antibody titres and protection in children aged 5-17 months in Kenya and Tanzania: a randomised controlled trial. Lancet Infect Dis 11: 102–109

Oyen D, Torres JL, Wille-Reece U, Ockenhouse CF, Emerling D, Glanville J, Volkmuth W, Flores-Garcia Y, Zavala F, Ward AB et al (2017) Structural basis for antibody recognition of the NANP repeats in Plasmodium falciparum circumsporozoite protein. Proc Natl Acad Sci U S A 114: E10438–E10445

Pompon J, Levashina EA (2015) A New Role of the Mosquito Complement-like Cascade in Male Fertility in Anopheles gambiae. PLoS Biol 13: e1002255

RTS Sctp, Agnandji ST, Lell B, Soulanoudjingar SS, Fernandes JF, Abossolo BP, Conzelmann C, Methogo BG, Doucka Y, Flamen A et al (2011) First results of phase 3 trial of RTS,S/AS01 malaria vaccine in African children. N Engl J Med 365: 1863–1875

Sattabongkot J, Yimamnuaychoke N, Leelaudomlipi S, Rasameesoraj M, Jenwithisuk R, Coleman RE, Udomsangpetch R, Cui L, Brewer TG (2006) Establishment of a human hepatocyte line that supports in vitro development of the exo-erythrocytic stages of the malaria parasites Plasmodium falciparum and P. vivax. Am J Trop Med Hyg 74: 708–715

Scally SW, Murugan R, Bosch A, Triller G, Costa G, Mordmuller B, Kremsner PG, Sim BKL, Hoffman SL, Levashina EA et al (2018) Rare PfCSP C-terminal antibodies induced by live sporozoite vaccination are ineffective against malaria infection. J Exp Med 215: 63–75

Stoute JA, Slaoui M, Heppner DG, Momin P, Kester KE, Desmons P, Wellde BT, Garcon N, Krzych U, Marchand M (1997) A preliminary evaluation of a recombinant circumsporozoite protein vaccine against Plasmodium falciparum malaria. RTS,S Malaria Vaccine Evaluation Group. N Engl J Med 336: 86–91

Swearingen KE, Lindner SE, Shi L, Shears MJ, Harupa A, Hopp CS, Vaughan AM, Springer TA, Moritz RL, Kappe SH et al (2016) Interrogating the Plasmodium Sporozoite Surface: Identification of Surface-Exposed Proteins and Demonstration of Glycosylation on CSP and TRAP by Mass Spectrometry-Based Proteomics. PLoS Pathog 12: e1005606

Tan J, Sack BK, Oyen D, Zenklusen I, Piccoli L, Barbieri S, Foglierini M, Fregni CS, Marcandalli J, Jongo S et al (2018) A public antibody lineage that potently inhibits malaria infection through dual binding to the circumsporozoite protein. Nat Med 24: 401–407

Thai E, Costa G, Weyrich A, Murugan R, Oyen D, Flores-Garcia Y, Prieto K, Bosch A, Valleriani A, Wu NC et al (2020) A high-affinity antibody against the CSP N-terminal domain lacks Plasmodium falciparum inhibitory activity. J Exp Med 217

Tiller T, Meffre E, Yurasov S, Tsuiji M, Nussenzweig MC, Wardemann H (2008) Efficient generation of monoclonal antibodies from single human B cells by single cell RT-PCR and expression vector cloning. J Immunol Methods 329: 112–124

Triller G, Scally SW, Costa G, Pissarev M, Kreschel C, Bosch A, Marois E, Sack BK, Murugan R, Salman AM et al (2017) Natural Parasite Exposure Induces Protective Human Anti-Malarial Antibodies. Immunity 47: 1197–1209 e1110

Wang LT, Pereira LS, Flores-Garcia Y, O’Connor J, Flynn BJ, Schon A, Hurlburt NK, Dillon M, Yang ASP, Fabra-Garcia A et al (2020) A Potent Anti-Malarial Human Monoclonal Antibody Targets Circumsporozoite Protein Minor Repeats and Neutralizes Sporozoites in the Liver. Immunity 53: 733–744 e738

Wang LT, Pereira LS, Kiyuka PK, Schon A, Kisalu NK, Vistein R, Dillon M, Bonilla BG, Molina-Cruz A, Barillas-Mury C et al (2021) Protective effects of combining monoclonal antibodies and vaccines against the Plasmodium falciparum circumsporozoite protein. PLoS Pathog 17: e1010133

Wang Q, Fujioka H, Nussenzweig V (2005) Mutational analysis of the GPI-anchor addition sequence from the circumsporozoite protein of Plasmodium. Cell Microbiol 7: 1616–1626

Wardemann H, Yurasov S, Schaefer A, Young JW, Meffre E, Nussenzweig MC (2003) Predominant autoantibody production by early human B cell precursors. Science 301: 1374–1377

WHO (2021) World malaria report 2021. Geneva: World Health Organization; 2021

Yaari G, Vander Heiden JA, Uduman M, Gadala-Maria D, Gupta N, Stern JN, O’Connor KC, Hafler DA, Laserson U, Vigneault F et al (2013) Models of somatic hypermutation targeting and substitution based on synonymous mutations from high-throughput immunoglobulin sequencing data. Front Immunol 4: 358

Zavala F, Cochrane AH, Nardin EH, Nussenzweig RS, Nussenzweig V (1983) Circumsporozoite proteins of malaria parasites contain a single immunodominant region with two or more identical epitopes. J Exp Med 157: 1947–1957

Zavala F, Tam JP, Hollingdale MR, Cochrane AH, Quakyi I, Nussenzweig RS, Nussenzweig V (1985) Rationale for development of a synthetic vaccine against Plasmodium falciparum malaria. Science 228: 1436–1440

